# Panpulmonate transcriptomes reveal candidate genes involved in the adaptation to freshwater and terrestrial habitats in gastropods

**DOI:** 10.1101/072389

**Authors:** Pedro E. Romero, Barbara Feldmeyer, Markus Pfenninger

## Abstract

**Background:** The conquest of the land from aquatic habitats is a fascinating evolutionary event that happened multiple times in different phyla. Mollusks are among the organisms that successfully invaded the non-marine realm, resulting in the radiation of terrestrial panpulmonate gastropods. We compared transcriptomes from panpulmonates to study the selective pressures that modeled the transitions from marine into freshwater and terrestrial realms in this molluscan lineage.

**Results:** *De novo* assembly of six panpulmonate transcriptomes resulted in 55,000 - 97,000 predicted open reading frames, of which 9 - 14% were functionally annotated. Adding published transcriptomes, we predicted 791 ortholog clusters shared among fifteen panpulmonate species, resulting in 702 amino acid and 736 codon-wise alignments. The branch-site test of positive selection applied to the codon-wise alignments showed twenty-eight genes under positive selection in the freshwater lineages and seven in the terrestrial lineages. Gene ontology categories of these candidate genes include actin assembly, transport of glucose, and the tyrosine metabolism in the terrestrial lineages; and, DNA repair, metabolism of xenobiotics, mitochondrial electron transport, and ribosome biogenesis in the freshwater lineages.

**Conclusions:** We identified candidate genes representing processes that may have played a key role during the water-to-land transition in Panpulmonata. These genes were involved in energy metabolism and gas-exchange surface development in the terrestrial lineages and in the response to the abiotic stress factors (UV radiation, osmotic pressure, xenobiotics) in the freshwater lineages. Our study expands the knowledge of possible adaptive signatures in genes and metabolic pathways related to the invasion of non-marine habitats in invertebrates.

## Background

The invasion from marine to non-marine habitats is one of the most enthralling events in the evolution of life on Earth. The transition from sea to freshwater and land environments occurred multiple times in different branches of the tree of life. Mollusks, along arthropods and vertebrates, are among the successful phyla that invaded the non-marine realm. Several branches from the molluscan class Gastropoda (Neritimorpha, Cyclophoroidea, Littorinoidea, Rissooidea, and Panpulmonata) have colonized terrestrial habitats multiple times [1, 2]. Especially, several independent land invasions in the Panpulmonata resulted in a significant adaptive radiation and explosive diversification that likely originated up to a third of the extant molluscan diversity [3]. Therefore, panpulmonate lineages are a promising system to study evolution of adaptations to non-marine habitats.

The habitat transition must have triggered several novel adaptations in behavior, breathing, excretion, locomotion, and osmotic and temperature regulation, to overcome problems that did not exist in the oceans such as dehydration, lack of buoyancy force, extreme temperature fluctuations and radiation damage [4-6]. Studies in vertebrates showed different genomic changes involved in the adaptation to the new habitats. Mudskippers, amphibious teleost fishes adapted to live on mudflats, possess unique immune genes to possibly counteract novel pathogens on land, and opsin genes for aerial vision and for enhancement of color vision [7]. Tetrapods showed adaptation signatures in the carbamoyl phosphate synthase I (CPS1) gene involved in the efficient production of hepatic urea [8]. Primitive sarcopterygians like the coelacanth *Latimeria* already possess various conserved non-coding elements (CNE) that enhance the development of limbs, and an expanded repertoire of genes related to the pheromone receptor VR1 that may have facilitated the adaptation to sense airborne chemicals during the water-to-land transition in tetrapods [9]. Also, vertebrate keratin genes responsible for skin rigidity underwent a functional diversification after the water-to-land transition, enhancing the protection against friction imposed by the new terrestrial lifestyle [10].

Conversely, information about the molecular basis of adaptation from marine to non-marine habitats in invertebrates is still scarce. Only one study reported adaptive signals in gene families (e. g. ATPases, DNA repair, and ribosomal proteins) that may have played a key role during terrestrialization in springtails and insects (Hexapoda) [11], clades that probably had a common pancrustacean ancestor living in a shallow marine environment [12, 13]. Mutations in the ATPases were suggested to provide the necessary energy to adapt to new high-energy demanding habitats [14], DNA repair genes would have helped reducing the damage produced by increased ultraviolet (UV) irradiation, and finally, as the ribosomal machinery is salt-sensitive, adaptive signs in the ribosomal proteins could have been a result of the different osmotic pressures within aquatic and terrestrial environments [15].

In a previous paper, we explored the adaptive signals in the mitochondrial genomes of panpulmonates [16]. We found that in the branches leading to lineages with terrestrial taxa (Ellobioidea and Stylommatophora), the mitochondrial genes *cob* and *nad5*, both involved in the oxidative phosphorylation pathway that finally produces ATP, appeared under positive selection. Moreover, the amino acid positions under selection have been related to an increased energy production probably linked to novel demands of locomotion [17, 18], and to changes in the equilibrium constant physicochemical property involved in the regulation of ROS production and thus, in the ability to tolerate new abiotic stress conditions [19].

Here, we expanded our search for candidate genes related to the adaptation to non-marine habitats, using transcriptome-wide data from several panpulmonate taxa, including marine, intertidal, freshwater and terrestrial lineages. We used a phylogenomic approach to reconstruct the evolutionary relationships of Panpulmonata and then tested for positive selection in the branches leading to freshwater and land snails. This approach aims to provide new insights into the selective pressures shaping the transition from marine to freshwater and land lifestyles.

## Results

We generated approximately 2,100,000 - 3,400,000 Illumina for our six samples (five ellobiids and one stylommatophoran species,Table 1). The quality trimming eliminated 14 - 39% of short and low-quality fragments in our samples. *De novo* meta assembly with MIRA produced approximately 55,000 - 98,000 transcripts in our samples and 54,000 - 130,000 in the other additional samples (Table 1). For further analyses we used transcripts larger than 300 bp. This represented a reduction of less than 1% in our samples but a higher reduction in the public data (3 - 35%). The number of predicted open reading frames was very similar to the number of transcripts > 300 bp in almost all cases, the only exception was *Radix balthica*, where only 57% of the transcripts obtained an ORF prediction. We obtained 9,000 - 30,000 single blast hits for our data, representing 5,000 - 13,000 single annotated genes. The percentage of annotated genes from our open reading frame data was 9 - 14%.

**Table 1.** Descriptive statistics of the assembled transcriptomes. *De novo* assemblies of new panpulmonate transcriptomes are highlighted in bold.

|  |  |  |  |  | Trimmed reads |  | MIRA meta-assembly |  |  |  | BlastX |  |
| --- | --- | --- | --- | --- | --- | --- | --- | --- | --- | --- | --- | --- |
| Clade | Species | Habitat | Accession<br>Number (SRA<br>database) | Number of<br>raw reads | For Trinity | For Bridger | All<br>transcripts | Transcripts ><br>300 (bp) | N50<br>(bp) | Predicted<br>ORF's | Single<br>hits | Single gene |
| Acochlidia | *Microhedyle*<br>*glandulifera* | Marine | SRR1505118 | 6194970 | 6103287 | 6028350 | 129453 | 84468 | 742 | 83983 | 28750 | 12772 |
| Acochlidia | *Strubellia wawrai* | Freshwater | SRR1505137 | 24132673 | 24049385 | 23436737 | 82681 | 79947 | 1582 | 79735 | 27911 | 13735 |
| Amphiboiloidea | *Phallomedusa solida* | Intertidal | SRR1505127 | 25685273 | 25496722 | 24822510 | 68633 | 65500 | 1424 | 64978 | 24560 | 12722 |
| **Ellobioidea** | ***Carychium* sp.** | **Terrestrial** | **XXXXXXXXXX** | **33608344** | **23461211** | **23461211** | **87994** | **87719** | **2035** | **87242** | **23586** | **11672** |
| **Ellobioidea** | ***Cassidula***<br>***plecotremata*** | **Intertidal** | **XXXXXXXXXX** | **26316221** | **18928297** | **17989294** | **62318** | **62031** | **1533** | **61569** | **9583** | **5717** |
| **Ellobioidea** | ***Melampus flavus*** | **Intertidal** | **SRR4103303** | **21068142** | **16919851** | **16052613** | **97629** | **97334** | **1640** | **96816** | **22187** | **10778** |
| Ellobioidea | *Ophicardelus sulcatus* | Intertidal | SRR1505124 | 16026272 | 15737826 | 15425623 | 74467 | 71073 | 1499 | 70573 | 20851 | 10937 |
| **Ellobioidea** | ***Pythia pachyodon*** | **Terrestrial** | **XXXXXXXXXX** | **24016251** | **20555474** | **19399330** | **92548** | **92264** | **2035** | **91759** | **24077** | **11776** |
| **Ellobioidea** | ***Trimusculus* sp.** | **Intertidal** | **SRR4102394** | **25613160** | **16343735** | **15696090** | **55728** | **55391** | **1240** | **55213** | **9310** | **5547** |
| Hygrophila | *Planorbarius corneus* | Freshwater | SRR1185333 | 28040804 | 27417761 | 26569686 | 86067 | 82815 | 2587 | 78934 | 39051 | 16959 |
| Hygrophila | *Biomphalaria glabrata* | Freshwater | SRR942795 | 172317158 | 170810180 | 164139487 | 70439 | 67493 | 2070 | 66689 | 33757 | 17015 |
| Hygrophila | *Radix balthica* | Freshwater | SRR097739 | 16923850 | NA | NA | 54450 | 54450 | 679 | 30883 | 14614 | 8140 |
| Pyramidelloidea | *Turbonilla* sp. | Marine | SRR1505139 | 26619896 | 26219791 | 25806515 | 132978 | 127301 | 1023 | 126409 | 34180 | 14316 |
| **Stylommatophora** | ***Arion vulgaris*** | **Terrestrial** | **XXXXXXXXXX** | **24874185** | **19179005** | **17977774** | **92316** | **91984** | **1454** | **91417** | **29208** | **13134** |
| Systellommatophora | *Onchidella floridana* | Intertidal | SRR1505123 | 14872953 | 14797399 | 14528352 | 79502 | 74540 | 1057 | 74146 | 28257 | 13679 |

We predicted 791 orthologous clusters shared among all species, of which 702 ortholog clusters remained after removing spurious and poorly amino acid aligned sequences in trimAL. From this dataset, MARE selected 382 informative clusters to reconstruct the phylogeny of the panpulmonate species (Additional File 1). The amount of missing data corresponds to 10.94% in the complete matrix, and 6.26% in the reduced matrix (Additional Files 2 and 3, respectively).

Most branches in the panpulmonate tree received high support (Figure 1). The clade containing Stylommatophora and Systellommatophora was significantly supported (bootstrap: 94 / posterior probability: 1.0) and appeared as a sister of the monophyletic Ellobioidea (99/1.0). The Acochlidia clade was moderately supported (86/1.0). The association of the Acochlidia with the Ellobioidea, Stylommatophora, Systellommatophora clade had no significant bootstrap support but a high posterior probability (64/1.0). The Hygrophila clade was highly supported (100/1.0). The association of Amphiboloidea and Pyramidelloidea was also highly supported (100/1.0).

We detected selection signatures on genes (codon-wise alignments) across the freshwater and terrestrial lineages in Panpulmonata. The likelihood-ratio test (LRT) comparing the branch-site model A against the null model (neutral) showed seven orthologous clusters under positive selection in the land lineages and twenty-eight clusters in the freshwater lineages (Additional File 4). There was no overlapping within positively selected genes from freshwater and terrestrial lineages. Table 2 shows examples of these candidate genes, their annotations, biological processes, molecular functions, and pathways involved. The BlastX annotations revealed candidate genes involved in the actin assembly, protein folding, transport of glucose, and vesicle transport in the terrestrial lineages. In the freshwater lineages, we found candidate genes associated to DNA repair, metabolism of xenobiotics, mitochondrial electron transport, protein folding, proteolysis, ribosome biogenesis, RNA processing and transport of lipids (Additional Files 5 and 6). We found no significant enriched GO (Gene ontology) terms neither in the freshwater nor terrestrial lineages.

**Table 2.** Examples of ortholog clusters under positive selection in the terrestrial and freshwater lineages. The complete information can be found in the Additional Files 4 and 5.

| Orthologous cluster | BlastX annotation | Molecular function | Biological process | KEGG pathway |
| --- | --- | --- | --- | --- |
| **Terrestrial** |  |  |  |  |
| OG0000060 | tyramine beta-hydroxylase-like | Copper ion binding, oxidoreductase activity | Oxidation-reduction process | Tyrosine metabolism |
| OG0000137 | sodium glucose cotransporter 4-like | Transmembrane transport | Transporter activity | Carbohydrate digestion and absorption |
| OG0001172 | alpha- sarcomeric-like isoform X2 | Actin filament binding, calcium ion binding | Actin crosslink formation, actin filament bundle assembly | Focal adhesion |
| **Freshwater** |  |  |  |  |
| OG0000120 | cytochrome P450 3A7-like | Monooxygenase activity, iron ion binding | Xenobiotic metabolic process | Aminobenzoate degradation, steroid hormone biosynthesis |
| OG0004116 | methyated-DNA-- -cysteine methyltransferase-like isoform X2 | methyated-DNA-[protein]-cysteine S methyltransferase activity | DNA repair | - |
| OG0004174 | cytochrome c oxidase subunit 4 isoform mitochondrial-like | Cytochrome c oxidase activity | Proton transport, mitochondrial electron transport, cytochrome c to oxygen | Oxidative phosphorylation |
| OG0002708 | 40S ribosomal protein S3a | - | RNA binding, protein binding | rRNA processing, translation |

Candidate genes under positive selection in the terrestrial lineages were involved in the carbohydrate digestion, endocytosis, focal adhesion, and the metabolism of lipids and tyrosine pathways. In case of the freshwater lineages, the candidate genes were involved in several metabolic pathways, for example, amino acid biosynthesis, focal adhesion, lysosome, oxidative phosphorylation, and protein signaling (Table 2, Additional Files 5 and 6).

## Discussion

Panpulmonates transitioned from marine to freshwater and terrestrial environments in several lineages and multiple times [20, 2, 21], Thus, they are a very suitable model to study the invasion of non-marine realms. However, the phylogenetic relationships within this clade are yet to be resolved [20]. Our tree topology using 382 orthologous clusters resembles the one obtained from Jörger et al. [21], based on mitochondrial and nuclear markers. In addition, we found support for the Geophila: Stylommatophora (terrestrial) and Systellommatophora (intertidal/terrestrial) as sister groups. This clade has been proposed before based on the position of the eyes at the tip of cephalic tentacles [22]. Still, previous phylogenies using mitochondrial and nuclear markers failed to support this clade [23, 16, 24, 21]. We also found support for Eupulmonata (*sensu* Morton [25, 23]), a clade comprising Stylommatophora and Systellommatophora plus Ellobioidea (intertidal/terrestrial) [20], this clade was supported using a combination of mitochondrial and nuclear markers [21]. Generation of high-quality transcriptomic data for other panpulmonate clades (marine Sacoglossa and Siphonarioidea, freshwater Glacidorboidea), and additional data for terrestrial Stylommatophora and Systellommatophora, will definitively illuminate the evolutionary relationships in Panpulmonata.

Our study is the first genome-wide report on the molecular basis of adaptation to non-marine habitats in panpulmonate gastropods. In case of the terrestrial lineages, we found evidence that the different positively selected genes are involved in a general pattern of adaptation to increased energy demands. The adaptive signs found in a gene related to actin assembly (OG0001172, Table 2) can be related to the necessity to move (forage, hunt preys or escape from predators) in the terrestrial realms. Moreover, the displacement in an environment lacking the buoyancy force to float or swim requires more energy, which can be obtained by increasing the glucose uptake (OG0000137) to produce energy in form of ATP. The adaptive signatures we found previously in two mitochondrial genes, *cob* and *nad5*, involved in energy production in the mitochondrion, also suggested a response to new metabolic requirements in the terrestrial realm, such as the increase of energy demands (to move and sustain the body mass).

One gene found under positive selection in the terrestrial genus *Pythia*, was involved in the metabolism of tyrosine (OG0000060). Tyrosine is the principal component of the thyroid hormones (TH). Despite invertebrates lack the thyroid gland responsible of the production of TH’s; the synthesis of TH’s has been demonstrated in mollusks and echinoderms. In these organisms, iodine is ligated to the tyrosine in the peroxisomes, producing thyroid hormones [26]. Notably, it has been suggested that iodinated tyrosine may have been essential in vertebrates during the transition to terrestrial habitats for TH’s are required in the expression of transcription factors involved in the embryonic development and differentiation of the lungs [27]. Land snails adapted to breath air by losing their gills and transforming the inner surface of their mantle into a lung [5]. Therefore, we propose that the tyrosine pathway was also a key component in invertebrates probably promoting the development of novel gas exchange tissues in land snails.

In case of the freshwater lineages, one of the positively selected genes was similar to the subunit 4 of the cytochrome c oxidase (*cob*) respiratory complex (OG0004174, Table 2). As mentioned above, *cob* is part of the energy production pathway in the cell. This enzyme complex contains many subunits encoded both in the mitochondrial and nuclear genome. The subunit 4 belongs to the nuclear genome and has an essential role in the assembly and function of the *cob* complex [28]. In agreement with our previous results that found the mitochondrial *cob* subunit under positive selection [16], we suggest that this gene was also involved in enhancing the metabolic performance of the enzyme and aided to cope with the new energy demands the realm transition.

A gene similar to cytochrome P450 was also found under positive selection (OG000120). Cytochrome P450s are proteins involved in the metabolism of xenobiotics. They were also under positive selection in the terrestrial Hexapoda lineages in comparison to other water-dwelling arthropods [11]. This result suggests that adaptations in these genes probably improve the response to new organic pollutants and toxins absent in the marine realm.

Another gene that showed adaptive signatures was the 40S ribosomal protein S3a (OG0002708). Likewise, ribosomal genes were also identified in a previous study on land-to-water transitions in hexapods [11] and plants [15]. In the latter study, it was suggested that the difference in the osmotic pressure from aquatic and terrestrial realms could affect the salt-sensitive ribosomal machinery, triggering adaptations to tolerate new salt conditions. This could also be the case for the freshwater animals (hypertonic) in comparison to the marine ones (hypotonic).

Finally, we found adaptive signatures in a DNA methyltransferase gene (OG0004116). This enzyme is part of the DNA repair system in the cell. Specifically, it removes methyl groups from O6-methylguanine produced by carcinogenic agents and it has been showed that its expression is regulated by the presence of ultraviolet B (UVB) radiation [29]. Positive selection on DNA repair genes has been found in hexapods [11], and in vertebrates living in high altitude environments (Tibetan antelopes) [14] or in mudflats (mudskippers) [7], suggesting an important role in the maintenance of the genomic integrity in response to the rise of temperature gradients or UV radiation in the terrestrial realms. In case of the aquatic environments, an extensive review has found an overall negative UVB effect on marine and freshwater animals [30]. However, the authors did not find a significant difference of the survival among taxonomic groups or levels of exposure in marine and freshwater realms, and suggested that the negative effects are highly variable among organisms and depends on several factors including cloudiness, ozone concentration, seasonality, topography, and behavior. Interestingly, it has been reported that survival in the freshwater snail *Physella acuta* (Hygrophila) depends of the combination of a photoenzymatic repair system plus photoprotection provided by the shell thickness and active selection of locations below the water surface avoiding the sunlight [31]

## Conclusions

We found that the positively selected genes in the terrestrial lineages were related to motility and to the development of novel gas-exchange tissues; while most of the genes in freshwater lineages were related to the response to abiotic stress such osmotic pressure, UV radiation and xenobiotics. These adaptations at the genomic level combined with novel responses in development and behavior probably facilitated the success during the transitions to the non-marine realm. Our results are very promising to understand the genomic basis of the adaptation during the sea-to-land transitions, and also highlight the necessity of more genome-wide studies especially in invertebrates, comparing marine, freshwater and terrestrial taxa, to unravel the evolution of the molecular pathways involved in the invasion of new realms.

## Methods

### Dataset collection

The dataset from Zapata et al. [32] was used as a starting point for our study. We added to this dataset the transcriptome from *Radix balthica* [33] and retrieved additional freshwater specimens from the NCBI Sequence Read Archive (SRA) (http://www.ncbi.nlm.nih.gov/sra). We complemented the dataset with five intertidal and terrestrial specimens from Ellobioidea (*Carychium* sp., *Cassidula plecotremata*, *Melampus flavus*, *Pythia pachyodon*, *Trimusculus sp*.) and one terrestrial Stylommatophora (*Arion vulgaris*), collected in Japan (2013) and Germany (2014), respectively (Additional File 7). RNA was isolated following the RNeasy kit (QIAGEN) following the manufacturer’s protocol. cDNA production and sequencing on the Illumina NextSeq500 platform (150 bp paired- end reads) was performed by StarSEQ GmbH (Mainz, Germany), according to their Illumina standard protocol. The final dataset comprised fifteen transcriptomes of panpulmonate species occurring in marine, intertidal, freshwater and terrestrial habitats (Table 1).

### Read processing and quality checking

FastQC [34] was used for initial assessment of reads quality. Then, Trimmomatic v0.33 [35] was used to remove and trim Illumina adaptor sequences and other reads with an average quality below 15 within a 4-base wide sliding window. In addition, we repeated the trimming analysis specifying a minimum length of 25 nt for further assembly comparisons. The same procedure was applied to all samples, except for *Radix* (454 reads). In this latter case, we got the transcriptome assembly directly from the author [33].

### Transcriptome assembly

*De novo* assembly was performed for all samples, except *Radix* (see last section), using Trinity v2.0.6 [36] with a minimum contig length of 100 amino acids, and Bridger v2014-12-01 [37] with default options. Bridger required the trimmed set with the minimum length of 25 nt. We combined the results from Trinity and Bridger in a meta-assembly using MIRA [38] with default settings. Only sequences with longer than 100 aa were retained for further analyses. This step was done to improve the accuracy in ortholog determination and facilitate phylogenomic analyses [39]. Furthermore, we used the ORFpredictor server [40] to predict open reading frames (ORF) within the transcripts.

### Construction of ortholog clusters

Ortholog clusters shared among protein sequences of the fifteen panpulmonate species were predicted using OrthoFinder [41] with default parameters. In case clusters contained more than one sequence per species, only a single sequence per species with the highest average similarity was selected using a homemade script. The predicted amino acid sequences from each ortholog cluster were aligned using MAFFT [42] with standard parameters. Nucleotide sequences in each orthogroup were aligned codon-wise using TranslatorX [43] taking into account the information from the amino acid alignments. Ambiguous aligned regions from the amino acid or codon alignments were removed using Gblocks [44] with standard settings. We used TrimAL [45] to remove poorly aligned or incomplete sequences in each ortholog cluster, using a minimum residue overlap score of 0.75.

### Phylogenomic analyses

Phylogenetic relationships among the Panpulmonata were reconstructed based on a subset of 382 ortholog clusters. The subset selection was done using MARE [46], a tool designed to find informative subsets of genes and taxa within a large phylogenetic dataset of amino acid sequences. The concatenated amino acid alignment length resulted in 88622 positions. Data were partitioned by gene using the partition scheme suggested in PartitionFinder [47] using the -*rcluster* option (relaxed hierarchical clustering algorithm), suitable for phylogenomic data [48]. We reconstructed an unrooted tree to be used as an input for the selection analyses. Maximum likelihood analyses were conducted in RAxML-HPC2 (8.0.9) [49]. We followed the “hard and slow way” suggestions indicated in the manual and selected the best-likelihood tree after 1000 independent runs. Then, branch support was evaluated using bootstrapping with 100 replicates, and confidence values were drawn in the best-scoring tree. Bayesian inference was conducted in MrBayes v3.2.2 [50]. Four simultaneous Monte Carlo Markov Chains (MCMC) were run, with the following parameters: eight chains of 20 million generations each, sampling every 20000 generations and a burn-in of 25%. Tracer 1.6 [51] was used to evaluate effective sample sizes (ESS > 200). We assume that a bootstrap value of >70% and a posterior probability of > 0.95 are evidence of significant nodal support.

### Selection analyses

The test of positive selection was performed for 736 ortholog clusters (codon-wise alignments) in CODEML implemented in the software PAML v4.8 [52]. PAML estimated the omega ratio (ω = dN: non-synonymous sites / dS: synonymous sites); ω = 1 indicates neutral evolution, ω < 1 purifying selection, and ω > 1 indicates positive selection [53]. To detect positive selection affecting sites along the terrestrial or freshwater branches (foreground) in comparison to the intertidal or marine lineages (background), the branch-site model A [54] in CODEML was applied (model = 2, NSsites = 2) for each orthologous cluster. The unrooted tree obtained using maximum likelihhod was set as the guide tree. In order to avoid problems in convergence in the log-likelihood calculations, we ran three replicates of model A with different initial omega values (ω = 0.5, ω = 1.0, ω = 5.0). We also calculated the likelihood of the null model (model = 2, NSsites = 2, fixed ω = 1.0). Both models were compared in a likelihood ratio test (LRT= 2*(lnL model A – lnL null model)). The Bayes Empirical Bayes (BEB) algorithm implemented in CODEML was used to calculate posterior probabilities of positive selected sites. We corrected p-values with a false discovery rate (FDR) cut-off value of 0.05 using the Benjamini and Hochberg method [55] implemented in R. The statistical significance of the overlap between positively selected genes from freshwater and terrestrial lineages was calculated using the R function *phyper*.

### Functional annotation

The transcripts were annotated using BlastX [56]. We blasted the nucleotide sequences against the invertebrate protein sequence RefSeq database (release 73, November 2015), with an e-value cutoff of 10^−6^. We selected top hits with the best alignment and the lowest e-value. Gene ontology (GO) terms for each BLASTx search were obtained in the Blast2GO suite [57]. Functional annotation information was obtained from InterPro database [58] using the InterProScan [59]. GO terms were then assigned to each orthologous group that was found under positive selection. In addition, we added to this clusters the metabolic pathway information retrieved from the KAAS server [60]. This server assigns orthology identifiers from the KEGG database (Kyoto Encyclopedia of Genes and Genomes). Functional enrichment analysis using the Fisher exact test was also performed in Blast2GO comparing the genes under positive selection against all ortholog clusters.

## Declarations

### Ethics approval and consent to participate

Not applicable.

### Consent to publish

Not applicable.

### Availability of data and materials

Raw sequence data is deposited in the Sequence Read Archive as BioProject (PRJNA339817), in the NCBI database. All other data sets (including trees, alignments, orthologous clusters, and scripts) supporting the results are available in the FigShare database: https://dx.doi.org/.

### Author’s contributions

PER carried out the fieldwork, transcriptome assembly, phylogenetic and molecular evolution analyses, conceived the study and wrote the manuscript. BF participated in the transcriptome assemblies and molecular evolution analyses, and helped to draft the manuscript. MP participated in the design and coordination of the study and revised the manuscript. All authors read and approved the final manuscript.

## Acknowledgements

We thank Yasunori Kano and Annette Klussmann-Kolb for their assistance in the fieldwork and Tilman Schell for his assistance in the bioinformatics analyses.

